# GALEON: A Comprehensive Bioinformatic Tool to Analyse and Visualise Gene Clusters in Complete Genomes

**DOI:** 10.1101/2024.04.15.589673

**Authors:** Vadim A. Pisarenco, Joel Vizueta, Julio Rozas

## Abstract

**Motivation:** Gene clusters, defined as a set of genes encoding functionally-related proteins, are abundant in eukaryotic genomes. Despite the increasing availability of chromosome-level genomes, the comprehensive analysis of gene family evolution remains largely unexplored, particularly for large and highly dynamic gene families or those including very recent family members. These challenges stem from limitations in genome assembly contiguity, particularly in repetitive regions such as large gene clusters. Recent advancements in sequencing technology, such as long reads and chromatin contact mapping, hold promise in addressing these challenges.

**Results:** To facilitate the identification, analysis, and visualisation of physically clustered gene family members within chromosome-level genomes, we introduce GALEON, a user-friendly bioinformatic tool. GALEON identifies gene clusters by studying the spatial distribution of pairwise physical distances among gene family members along with the genome-wide gene density. The pipeline also enables the simultaneous analysis and comparison of two gene families, and allows the exploration of the relationship between physical and evolutionary distances. This tool offers a novel approach for studying the origin and evolution of gene families.

**Availability and Implementation:** GALEON is freely available from http://www.ub.edu/softevol/galeon, and from https://github.com/molevol-ub/galeon

## 1. Introduction

Gene clusters, which encompass sets of genes encoding functionally-related proteins, are common in eukaryotic genomes. One prevalent type of gene cluster is the gene family, which comprises homologous genes generated from gene duplication, often facilitated by unequal crossing-over, leading to their arrangement in tandem within the genome (Ohno, 1970, Leister, 2004, Legan et al. 2021). However, despite the increasing availability of complete genome assemblies for many species in recent years (Ellegren, 2014, Bleidorn, 2016, Michael and VanBuren 2020), the comprehensive evolutionary analysis of gene family members remains largely unexplored. This results from several limitations, including: i) DNA sequencing errors introduced by the sequencing technologies, ii) incomplete and fragmented genome assemblies caused by repetitive elements, iii) inaccurate genome annotation. All these limitations have hindered the fine-scale analysis of medium-sized gene families (from ∼10 to ∼100 members), while large-sized families (comprising hundreds or thousands of members) are usually beyond the capabilities of current technologies. However, with the decreasing sequencing costs, advancements in high-quality long-read sequencing technologies (Hon et al. 2020, De Coster et al. 2021, Bi et al. 2024), and the development of specific bioinformatic tools for annotating gene family members in genome-wide data, such as BITACORA (Vizueta et al. 2020a, 2020b) or InsectOR (Karpe et al. 2021), there is promising potential to address these limitations.

Two critical challenges have precluded such comprehensive analysis: large gene family size and the presence of very recent (young) members. These aspects are problematic because of the limitations of current genome assemblies, even those employing long reads, which cannot accurately assemble large stretches of repetitive DNA regions encompassing highly diverged and recently originated copies (Vieira, et al. 2007, Librado and Rozas, 2013, Clifton et al. 2020). These limitations compromise the accurate estimation of rates for the origin of members through gene duplication (via exchange of DNA fragments between tandemly arranged repeats) and those of gene conversion likely leading to underestimation. As a result, it precludes a fine analysis of the impact of gene conversion versus an independent divergence in gene family evolution (Nei and Rooney, 2005, Eirin-Lopez et al. 2012). This fact has relevant implications; if gene conversion were more ubiquitous in gene family evolution, the inference of phylogenetic relationships could be misleading. The increasing number of chromosome-resolved assemblies addresses these limitations, allowing comprehensive analyses to understand gene family evolution and its genomic organisation.

Despite the availability of some bioinformatic tools like ClusterScan (Volpe et al. 2018) and C-Hunter (Yi et al. 2007), which were designed as unbiased methods for finding clusters, there is currently no comprehensive suite that integrates these analyses using genome-wide (chromosome-level) data, while also provide functional insights and helpful visualisation tools. To overcome such limitations, we introduce GALEON, a user-friendly bioinformatic tool designed to identify, analyse and visualise physically clustered gene family members in chromosome-level genomes. GALEON uses simple input file formats with gene coordinates (GFF3 or BED) and (optionally) protein sequence data. Specifically, the software assesses the cluster organisation of a given gene family by analysing the distribution of pairwise physical distances between genes, taking into account the average gene density across the genome. Moreover, if protein information is provided, GALEON can also be used to analyse the relationship between physical and evolutionary distances, providing insights into the origin and evolution of gene family members. Finally, GALEON also allows the simultaneous study of two gene families at once to explore putative co-evolving gene families.

## 2. Methods and implementation

GALEON implements an algorithm to identify gene family clusters in genome assemblies, analyse the distribution of physical and genetic distances, and present the results in tables and plots, which are then summarised in an HTML report (Figure 1). Depending on the type of analysis conducted, GALEON requires of the following data and input files: i) the genome size (in Mb); ii) the coordinate file containing the focal (one or two) gene family members in BED or GFF3 format; iii) the proteins encoded by the gene family members included in the coordinates file in FASTA format. The software is written in Python, bash and R.

**Figure 1.**
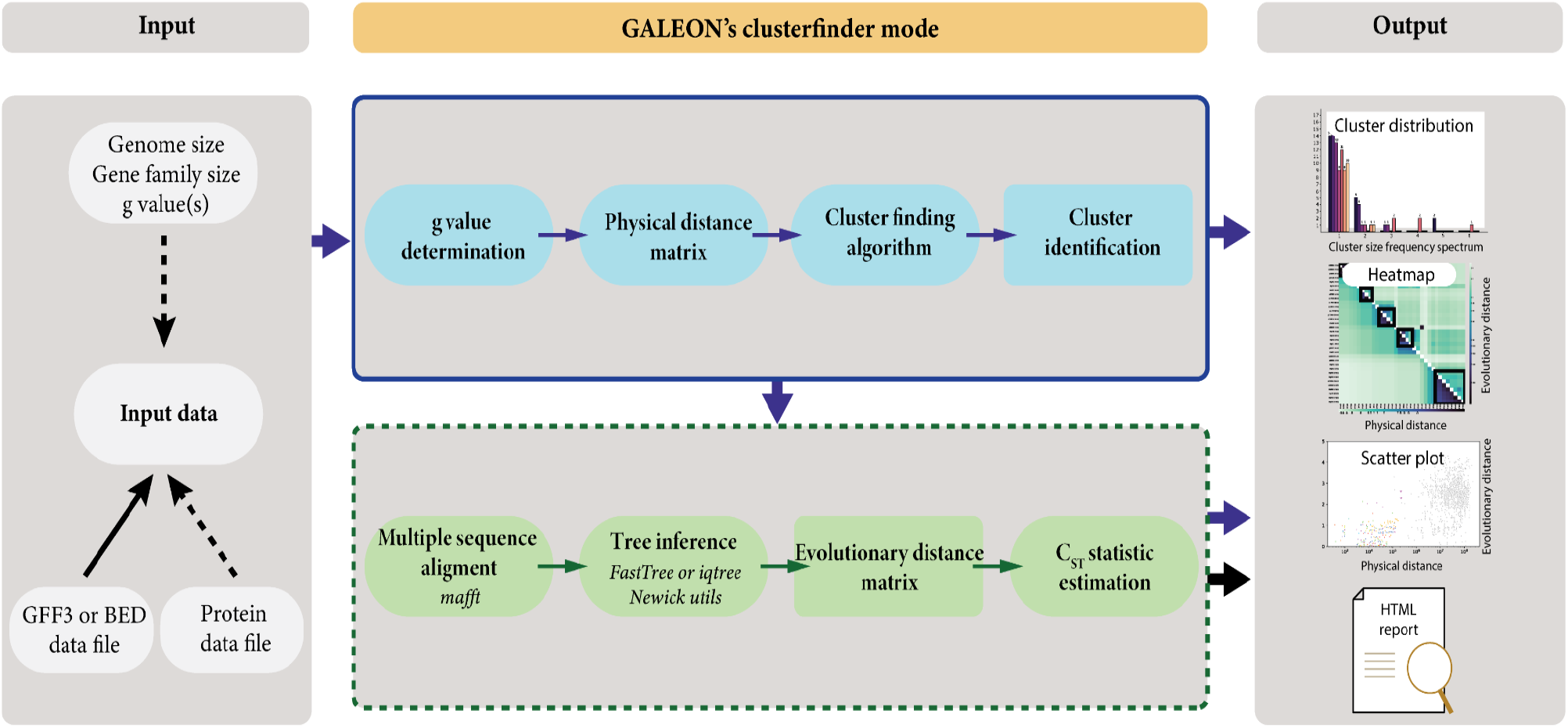
Schematic representation of the GALEON workflow. Solid lines represent mandatory data and steps, while dashed lines denote optional ones.

### 2.1 Gene cluster identification

GALEON identifies gene family clusters by analysing whether paralogous members are physically closer than expected by chance given the genome density of gene family members. In this context, we consider that *n* physically close members are clustered if they are arranged within a genomic region spanning less than specified cut-off *C*_*L*_ value (Vieira et al. 2007, Escuer et al. 2022):

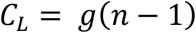

where *C*_*L*_ is the maximum length of a cluster containing two or more copies of the same family, while *g* is the maximum physical distance between two members to be considered as clustered. The *p*-values associated with a given *g* value are estimated under the assumption that the number of members follows a Poisson distribution. While the *g* values are specific to a particular gene family, selecting a lower *p*-value will prevent the inclusion of false positives in the majority of cases.

### 2.2 Gene cluster analyses

GALEON performs evolutionary analyses to provide insights into the origin, maintenance, and fate of gene family members, incorporating a comparative analysis of physical and evolutionary (genetic) distances. First, MAFFT (Katoh and Standley, 2013) is used to build a protein multiple sequence alignment (MSA) encompassing all gene family members. Subsequently, either FastTree (Price et al. 2009) or IQ-TREE (Minh et al. 2020) is employed to reconstruct the phylogeny of the gene family using the generated MSA. FastTree, employing default parameters under the JTT model, offers significantly faster computation compared to IQ-TREE, which uses ModelFinder to select the best substitution model (Kalyaanamoorthy et al. 2017). However, the enhanced accuracy of IQ-TREE’s phylogeny comes at the expense of a longer computational time. The resulting tree is used to estimate evolutionary distances, measured as the number of amino acid replacements per amino acid site, across all pairwise comparisons. The resulting distance matrices serve as the basis for exploring the relationship between physical and evolutionary distances. GALEON implements the *C*_*ST*_ statistic (Escuer et al. 2022), which measures the proportion of the genetic distance attributable to unclustered genes:

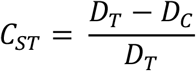

where *D*_*T*_ is the average of pairwise distances between all gene family copies, while DD_CC_ denotes the average of pairwise distances within a cluster. GALEON uses the Mann– Whitney U-test to determine whether the evolutionary distances within genomic clusters are significantly different from those of unclustered members. *C*_*ST*_ values are estimated separately for each chromosome (or scaffold), as well as for the whole genome data.

### 2.3 Output and Visualisation

GALEON results consist of several tables and figures that are also presented in a HTML report (Figure 2). Summary tables provide an overview of the general organisation of the focal gene family, categorised as “clustered” or “singleton” (not clustered members). Additionally, more detailed tables offer expanded information on the gene cluster members. Bars plots illustrate the gene cluster frequency spectrum, or distribution of the cluster sizes across scaffolds or the entire genome, while heatmaps depict the location of every gene cluster on each scaffold. If the corresponding protein data files are provided, GALEON generates additional heatmaps and scatter plots, showing physical distances plotted against evolutionary distances. Furthermore, GALEON’s parameters can be easily adjusted to customise the output plots according to user preferences.

**Figure 2.**
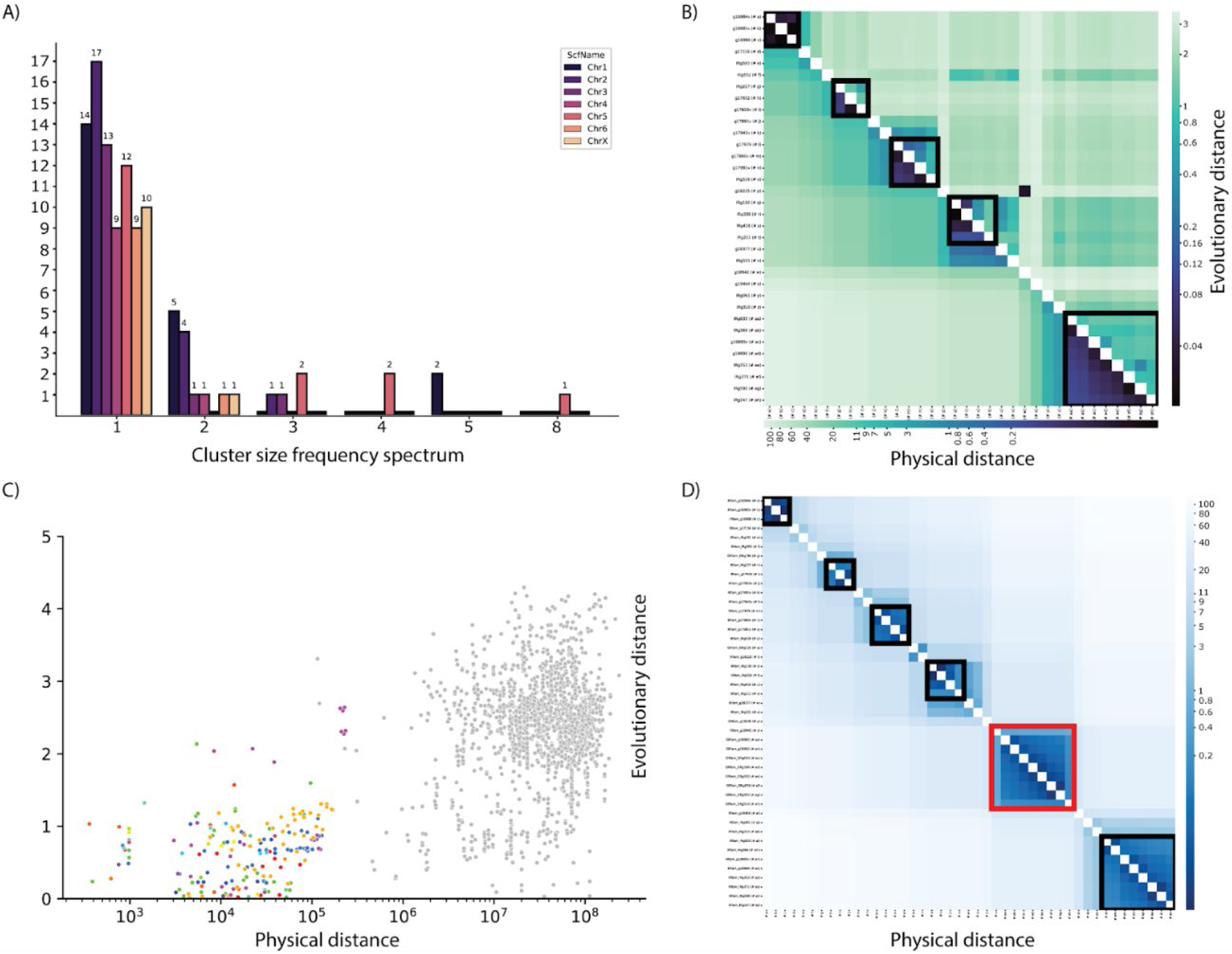
Overview of various visualisation tools and features integrated in GALEON, using data from the ionotropic (IR) gene family reported in Escuer et al. (2022). A) Gene cluster size frequency spectrum distribution across different chromosomes (colour-coded). The x-axis represents cluster size, while the y-axis represents the frequency. B) Heatmap showing the comparison of physical distances (lower triangular matrix; in Mb units) and evolutionary distances (upper triangular matrix; in amino acid substitutions per site units) in the chromosome 5 of *Dysdera silvatica*. The identified clusters are enclosed within black square shapes. C) Scatter plot depicting physical vs. evolutionary distances. Distances between clustered genes are coloured, while those of singletons are shown in grey. D) Heatmap displaying the joint analysis of the IRs and GRs gene families in the chromosome 5 of *Dysdera silvatica*. Black squares highlight clusters formed by members of one family, while red squares highlight those formed by members of both families.

## 3. Conclusions

We have developed a comprehensive bioinformatic tool designed to facilitate the identification, analysis, and visualisation of physically clustered gene families in chromosome-level genomes. Overall, GALEON offers a novel tool for studying clustered genes, by facilitating integrated analysis of physical and evolutionary distances, as well as the exploration of gene family co-evolution. Therefore, GALEON serves as a valuable starting point for gaining insights into the origin, maintenance, functional significance, and evolution of gene families.

## Acknowledgments

We thank all beta testers, whose feedback helped us to significantly improve the software, especially all people from the Evolutionary Genomics & Bioinformatics group at the Universitat de Barcelona.

## Funding

This work was supported by the Ministerio de Ciencia e Innovación of Spain (MCIN/AEI/10.13039/501100011033; grants PID2019-103947GB-C21 and PID2022-138477NB-C22 to J.R.; FPIs fellowships PRE2020-095592 to V.A.P), and from Comissió Interdepartamental de Recerca I Innovació Tecnològica of Catalonia, Spain (2021SGR00279).

## Notes

https://github.com/molevol-ub/galeon

### Competing Interest Statement

The authors have declared no competing interest.

